# Joint protein, mRNA and DNA sequence design and optimization with nucleotide-level Potts models

**DOI:** 10.64898/2026.10.05.756407

**Authors:** Błażej Banaszewski, Lars J. Dornfeld, Dexiong Chen, Karsten Borgwardt, Lukas F. Milles

**Affiliations:** Department of Machine Learning and Systems Biology, Max Planck Institute of Biochemistry, Martinsried, Germany; Research Group Biomolecular Design, Max Planck Institute of Biochemistry, Martinsried, Germany; Gene Center and Department of Biochemistry, LMU Munich, Munich, Germany

## Abstract

Protein design typically generates amino acid sequences first and then translates them into DNA post-hoc limiting control over the nucleotide sequence via codon choice at design time. We introduce NuCaliby, a structure-conditioned model that jointly designs amino acid, mRNA, and DNA sequences of a protein at nucleotide resolution. NuCaliby predicts a Potts model over a nucleotide graph, trained using amino acid supervision by marginalizing over synonymous codons. It preserves residue-level designability on par with the original Caliby model. Its energy-based formulation supports composable inference-time guidance from sequence-level objectives through proposal construction and candidate acceptance, allowing the incorporation of biological proxies into the coding nucleotide sequence. Under organism-specific tRNA adaptation guidance, NuCaliby improves adaptation to host tRNA pools beyond synonymous sequence space, trading-off self-consistency. NuCaliby also embeds specified RNA motif sequences directly into coding sequences, avoiding the combinatorial growth required for the same task at the amino-acid level. In wet lab experiments, NuCaliby produces expressible and soluble proteins on par with state of the art methods, while enabling more complex nucleotide-level control and optimization. Together, these results extend protein design beyond the amino acid level, enabling joint optimization across the central dogma of molecular biology. The code and model weights are available here: github.com/blazejba/NuCaliby

## 1 Introduction

Protein design is commonly formulated as a two-stage process: backbone generation followed by sequence design through inverse folding [7, 13, 1, 12]. Existing inverse folding approaches operate at the residue level, predicting amino acid sequences that are assumed to fold into the target structure. Here, we introduce NuCaliby, a nucleotide-level inverse folding framework that jointly designs protein, mRNA, and DNA sequences under composable nucleotide-level constraints beyond synonymous space.

In nature, protein expression follows the central dogma of molecular biology [6]: a DNA sequence encoding a design is introduced into a host organism, such as *E. coli*, transcribed into mRNA, and translated into the corresponding protein sequence. This introduces an additional layer of complexity, as the mapping from codons to amino acids is degenerate, with multiple synonymous codons encoding the same residue. While this degeneracy expands the nominal sequence space, synonymous codon choices are not necessarily neutral. They influence translation efficiency, mRNA stability, and other regulatory properties, and are often adapted differently across organisms and contexts [10, 26, 42, 43].

As a result, the set of nucleotide sequences compatible with a given protein in its biological context is heavily constrained in practice. Biological systems provide clear evidence of such constraints: for example viral genomes are under strong pressure to minimize their size and therefore frequently encode multiple functional elements within the same sequence, including regulatory RNA structures and even overlapping protein-coding sequences, by using the flexibility provided by the degenerate codons. [9].

**Figure 1.**
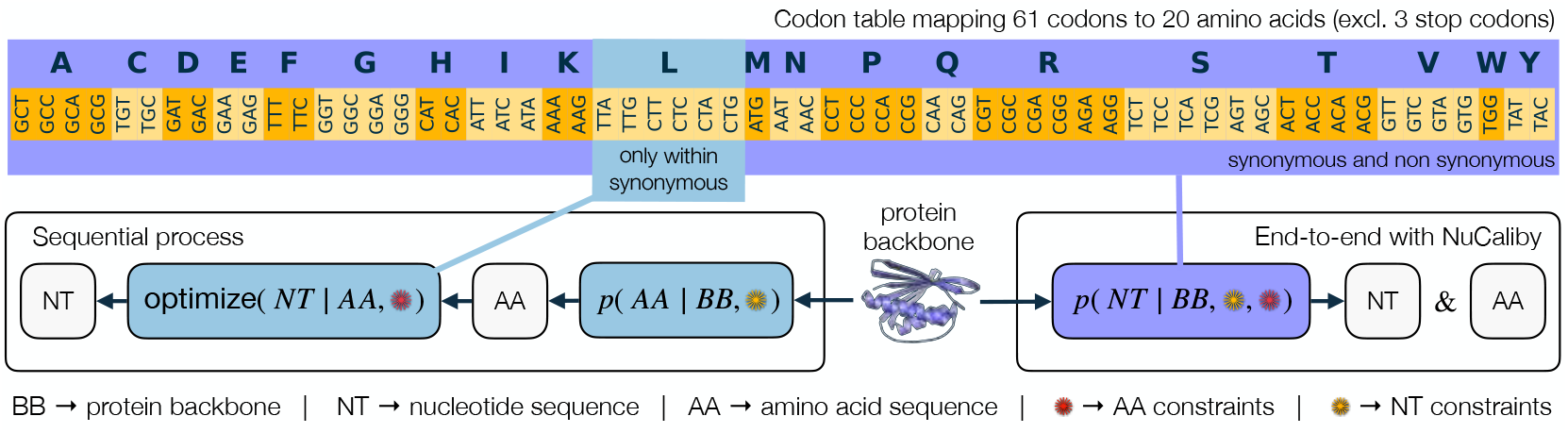
Conceptual differences between protein design performed sequentially on amino acid level and end-to-end on nucleotide-level. The former determines the amino acid sequence first, and optimization for DNA and RNA properties happens post hoc with optimization space limited to synonymous codons. NuCaliby enables amino acid and nucleotide level constraints to be enforced jointly during sampling, which allows design and optimization across the central dogma of life.

Current protein design pipelines typically choose an amino-acid sequence first and optimize its DNA encoding afterward. Yet many different amino-acid sequences can fold into the same backbone, and each has many DNA encodings [32]. Choosing the protein sequence first therefore limits the options available for satisfying DNA- and RNA-level objectives. NuCaliby keeps both choices open, bringing sequence design across the central dogma into a single optimization process. Methods that bridge this gap, such as CodonMPNN [28], predict codons directly from backbone but inherit average codon-usage statistics from natural genes and offer no inference-time control over downstream properties such as expression or mRNA stability.

Beyond this decoupling, existing inverse folding models [7, 28, 27] share an architectural limitation: they operate on residue-level graphs whose nodes represent amino acids or codons, obscuring nucleotide-level interactions such as dependencies across codon boundaries and long-range base-pairing effects that govern mRNA secondary structure.

These limitations motivate a finer-grained modeling approach that operates directly at nucleotide resolution. We introduce NuCaliby, a structure-conditioned Potts framework that co-designs protein and DNA sequences in a single pass. NuCaliby predicts both residue-level and nucleotide-level Potts parameters from the input protein structure, enabling sampling at either level. Crucially, the energy-based formulation provides a natural mechanism for incorporating external guidance potentials at inference time, analogous to classifier guidance in diffusion models, without retraining [37].

These energy biases allow navigating the multi-objective landscape of protein and nucleotide properties at generation time. By operating at nucleotide resolution, properties that were previously fixed at reverse-translation or ignored entirely, such as codon adaptation, mRNA stability, and RNA sequence motif inclusion become explicit yet adaptable design objectives alongside protein sequence-structure compatibility.

Our main contributions are: (a) a nucleotide-level Potts model for joint protein, mRNA, and DNA sequence design, enabling simultaneous optimization across the central dogma, (b) an inference-time energy guidance framework that allows controllable multi-objective optimization, e.g. tRNA adaptation, RNA motif incorporation or improvement in calculated RNA-stability, without retraining, (c) computational and wet lab experimental evidence that nucleotide-level control can be introduced with minimal loss in protein designability, while enabling objectives that are difficult or infeasible in synonymous sequence space.

## 2 Related work

### Inverse folding

Inverse folding models take a target backbone as input and generate sequences predicted to fold into that structure. ProteinMPNN [7], building on earlier work by John Ingraham *et al*. [12], remains the most widely used protein design method [38]. It is a graphical model based on message-passing neural networks, where residues are represented as nodes, while edges are featurized using pairwise geometric relationships derived from local protein coordinate frames, using autoregressive sampling for sequence generation. Chroma is among the first methods to use neural-network-derived Potts models for inverse folding [18, 13]. Neural Potts models represent functional energy landscapes of protein sequence alignment to a given structure. Caliby [27], an extension of neural Potts models, introduces protein-ensemble-based averaging of the Potts parameters using ensembles generated by partial diffusion with Protpardelle [5]. With this Caliby partially removes non-structural biases tied to a single backbone, and shows consistent improvements in in silico designability metrics (self-consistency and pLDDT).

### Codon optimization

Designed amino-acid sequences are typically reverse-translated post-hoc. DNA Chisel [43] applies rule-based constraints (codon adaptation, restriction-site avoidance, GC content); deep-learning approaches [40] map amino-acid to nucleotide sequences within the synonymous space using patterns from natural sequences. CodonMPNN [28] instead learns codon preferences end-to-end from genomic sequences paired with AlphaFold-predicted structures, conditioning on organism labels. Organism conditioning, however, requires taxonomically diverse training data that PDB alone cannot provide; the resulting reliance on AFDB structures degrades inverse-folding designability [31] (Appendix F), and natural genomic sequences are not necessarily optimized for expression or synthesizability.

### Inference-time guidance for discrete sampling

Classifier guidance has been extended to discrete sequence generation and applied to autoregressive inverse-folding models [37, 24]. During sampling, an auxiliary classifier biases transition probabilities toward sequences with desired properties, either by explicitly evaluating all single-mutant candidates at each step or by using a first-order Taylor approximation to adjust transition probabilities using classifier gradients. In both cases, the guiding classifier must accept partial or masked sequences, since autoregressive generation produces sequences one token at a time. Neural-Potts inverse-folding approaches such as Caliby [27] use discrete MCMC to sample complete sequences. This provides a natural setting for objectives that require fully specified sequences, motivating our use of DLMC [30] for nucleotide-level guidance.

## 3 Inverse folding with nucleotide-level Potts models

### Problem statement

Given a protein backbone *X* over *N* residues, inverse folding seeks a sequence that encodes a protein folding into *X*. *Amino-acid design* predicts a residue sequence *a* = (*a*_1_, …, *a*_*N*_) *∈ A*^*N*^ over the amino-acid alphabet *A* with |*A*| = 20. *Nucleotide-level design* predicts a coding sequence 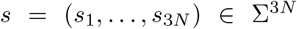 over the nucleotide alphabet Σ = *{A, C, G, T}*, with amino acids recovered through the genetic code *τ* : Σ^3^ *→ A*. We write syn(*a*) = *τ*^*−*1^(*{a}*) *⊆* Σ^3^ for the codons translating to a single amino acid *a*.

Classical codon optimisation first fixes an amino-acid target *a*^*⋆*^ *∈ A*^*N*^ and searches only within its synonymous codon space 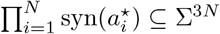, optimising an auxiliary objective *E*_bias_ subject to a hard synonymous constraint. NuCaliby instead models the full nucleotide space Σ^3*N*^ with a structure-conditioned Potts: amino acid sequence is encoded as a soft preference by the model rather than being enforced. Sampling can therefore move outside this synonymous space for any single *a*^*⋆*^ when this improves the trade-off between designability and *E*_bias_.

A Potts model on a graph *G* = (*V, ℰ*) over an alphabet Ω assigns each configuration *x ∈* Ω^|*V*|^ the energy

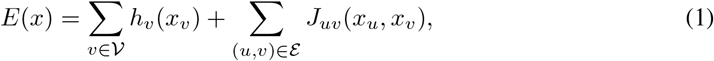

with single-site fields *h*_*v*_ and pairwise couplings *J*_*uv*_ predicted by the model. The induced distribution *p*_Potts_(*x*) *∝* exp(*−E*(*x*)) has a partition function over Ω^|*V*|^ that is intractable in general, so we never form it. Training and sampling rely only on site conditionals

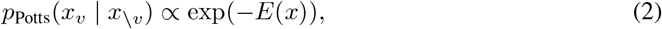

in which the partition function cancels and the per-site normaliser sums over Ω alone. NuCaliby predicts two such models conditioned on the backbone: a residue-level Potts on *G*^res^ over *A* and a nucleotide-level Potts on the expanded *G*^nuc^ over Σ.

NuCaliby is trained as a discrete masked denoiser. At each step we sample a conditioning level *t ∈* [0, 1], the probability of retaining a residue’s sequence identity in the model input. Residues are independently masked with probability 1 *− t*, with at least one residue masked per example; all three nucleotide positions of a masked codon are masked together. Thus, high *t* corresponds to mostly visible sequence and low *t* to heavily masked sequence. The model reconstructs native tokens from the backbone *X* and corrupted sequence 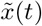 at inference we sample from the learned Potts model with optional auxiliary guidance.

### 3.1 Architecture and nucleotide graph construction

NuCaliby’s encoder, an AtomMPNN [27], processes the input residue graph constructed from protein structure into node and edge embeddings, which then split into two parallel decoder pathways predicting residue-level and nucleotide-level Potts parameters respectively (Figure 2).

**Figure 2.**
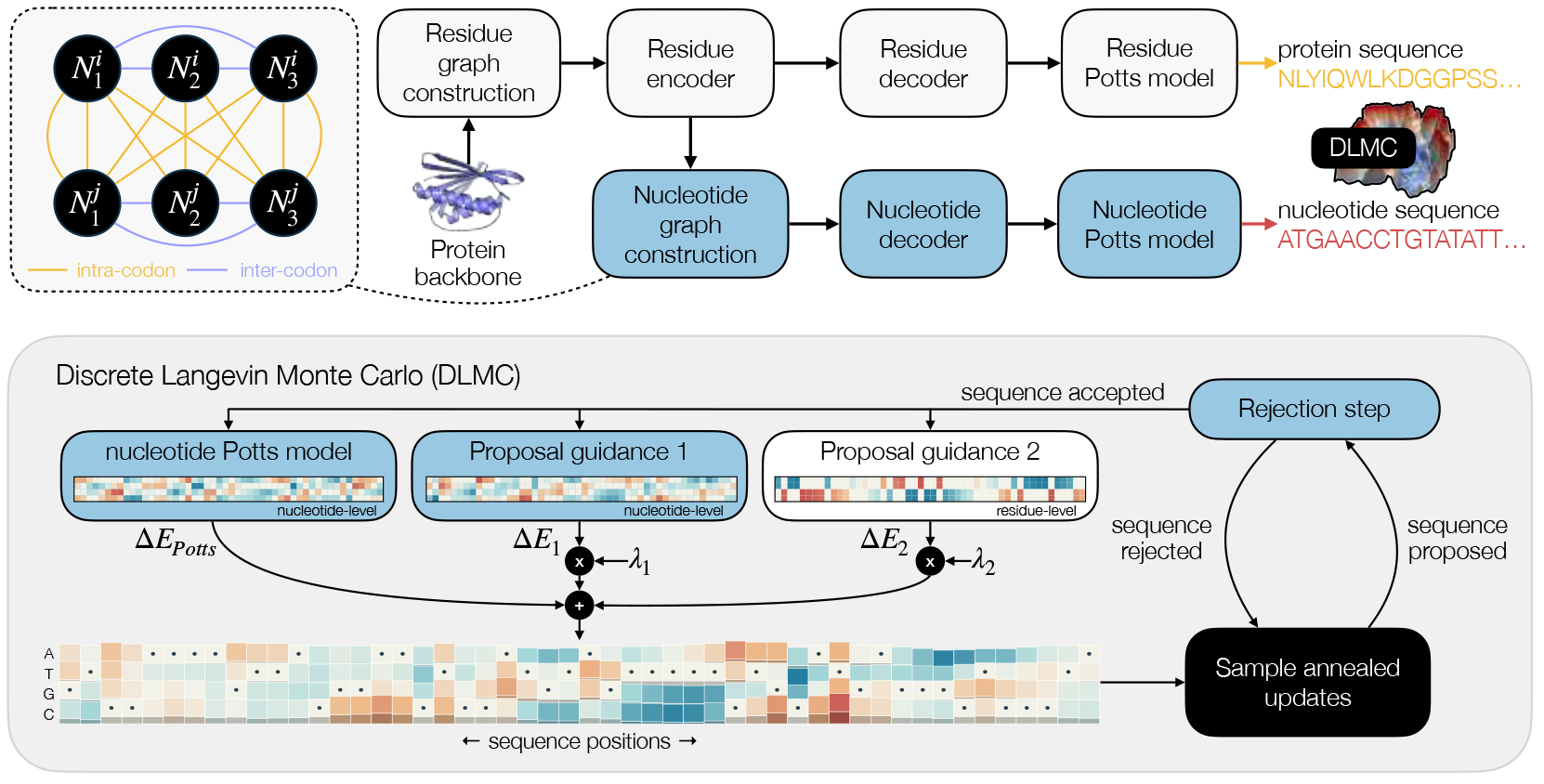
NuCaliby predicts the parameters of both residue- and nucleotide-level Potts models. The nucleotide graph is constructed with intra- and inter-codon edges explicitly distinguished, helping the model learn both codon-level organisation and structure-sequence relationships. DLMC explores the sequence space by inducing an energy field around the current sequence to satisfy structural compatibility, measured by the Potts model, and other objectives, which enter the optimization via proposal guidance and rejection step.

Given a backbone with *N* residues, we build a residue-level *k*-NN graph with *N* nodes and *Nk* directed edges (including self-loops). We lift it to a nucleotide graph with 3*N* nodes and 3*N* (3*k −* 1) directed edges. Intra-codon edges allow direct interactions among the three nucleotides encoding a residue, while inter-codon edges connect nucleotide choices across structurally neighbouring residues. This preserves codon organization while allowing dependencies between codons; Table 2 assesses the contribution of each edge family. Let 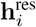, 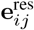 be the residue-level node and edge embeddings (with *i, j ∈ {*1, …, *N}* indexing residues), and 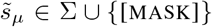 the (possibly masked) nucleotide input at position *µ ∈ {*1, …, 3*N}*. Nucleotide nodes and edges are featurised as

**Table 1:** Fraction of recovered amino acids (AA %), prediction confidence *(pLDDT* %), and self-consistency TMscore and RMSD in Å.

| | AA ↑ | pLDDT ↑ | TM ↑ | RMSD ↓ |
| --- | --- | --- | --- | --- |
| NATIVE | 100 | 40.9 ± 14.8 | .372 ± .166 | 5.21 ± 1.54 |
| PROTEINMPNN | 44.4 ± 7.7 | 54.3 ± 18.7 | .509 ± .233 | 4.37 ± 1.81 |
| CODONMPNN | 44.5 ± 9.0 | 47.0 ± 17.1 | .441 ± .211 | 4.85 ± 1.75 |
| CALIBY | 39.2 ± 7.2 | 61.7 ± 19.4 | .597 ± .248 | 3.85 ± 1.88 |
| CALIBY<sub>ENS</sub> | 28.5 ± 5.9 | 63.8 ± 16.8 | .591 ± .227 | 3.82 ± 1.71 |
| NUCALIBY<sup>AA</sup> | 39.6 ± 7.2 | 61.8 ± 19.6 | .593 ± .245 | 3.84 ± 1.85 |
| NUCALIBY<sup>AA</sup><sub>ENS</sub> | 29.2 ± 5.3 | 67.7 ± 17.2 | .659 ± .229 | 3.48 ± 1.75 |
| NUCALIBY<sup>NT</sup> | 38.6 ± 7.1 | 60.8 ± 19.1 | .579 ± .245 | 3.85 ± 1.82 |
| NUCALIBY<sup>NT</sup><sub>ENS</sub> | 27.2 ± 6.0 | 63.6 ± 16.3 | .580 ± .228 | 3.92 ± 1.64 |

**Table 2:** Ablations of nucleotide-level designability. Mean *±* standard deviation across 183 test proteins; RMSD in Å.

| NT variant | pLDDT ↑ | TM ↑ | RMSD ↓ |
| --- | --- | --- | --- |
| FULL MODEL | 60.8 ± 19.1 | .579 ± .245 | 3.85 ± 1.82 |
| NO INTRA-CODON | 55.1 ± 18.2 | .504 ± .225 | 4.40 ± 1.72 |
| NO INTER-CODON | 54.1 ± 17.7 | .500 ± .223 | 4.49 ± 1.80 |
| NT-ONLY TRAINING | 58.9 ± 18.5 | .548 ± .227 | 4.15 ± 1.73 |

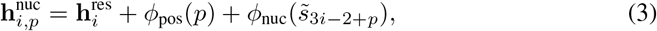

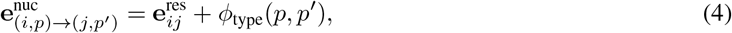

where *p, p*′ *∈ {*0, 1, 2*}* index codon positions and *ϕ*_pos_, *ϕ*_nuc_, *ϕ*_type_ are learnable embeddings; intracodon edges (*i* = *j*) set 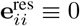 since no residue-level edge exists.

The nucleotide decoder refines the node and edge features of the nucleotide graph, which are then projected to predict Potts fields 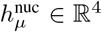 at each nucleotide site *µ ∈ {*1, …, 3*N}* and couplings **J**_*µν*_ *∈* ℝ^4*×*4^ for all edges (*µ, ν*) *∈ ℰ*^nuc^. We use a factorized parameterization **J**_*µν*_ = **L**_*µν*_**R**_*µν*_, where the matrices **L**_*µν*_, **R**_*µν*_ *∈* ℝ^4*×*4^ are predicted from the refined edge features.

### 3.2 Training

We optimise a joint objective combining residue-level and nucleotide-level losses, training the entire model end-to-end from scratch. The residue-level objective provides an auxiliary supervision signal for the shared structural encoder. Joint training modestly improves nucleotide-level designability compared with NT-only training, while residue-level performance remains similar (Appendix E). Organism-specific codon preferences are not learned from genomic data; they are introduced at inference through tAI-based energy biases that proxy translation efficiency (§3.3).

Write *c*_*i*_ = (*s*_3*i−*2_, *s*_3*i−*1_, *s*_3*i*_) *∈* Σ^3^ for the codon at residue *i*. Alongside the Potts head, the decoder outputs a per-position categorical 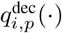 over Σ for each codon position *p ∈ {*0, 1, 2*}*, giving a factorised codon distribution 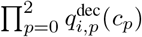 for *c* = (*c*_0_, *c*_1_, *c*_2_).

At each training step we sample a synonymous nucleotide sequence *s* ~ П_*i*_ syn(*a*_*i*_) from the native amino-acid sequence. This *s* plays two roles: a corrupted copy 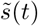 feeds the model as input, while the uncorrupted 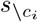 supplies the conditioning context for the Potts conditional. Because the training target specifies an amino-acid sequence rather than a preferred nucleotide encoding, supervision marginalises over the synonymous codon set instead of treating the sampled encoding as the unique correct target:

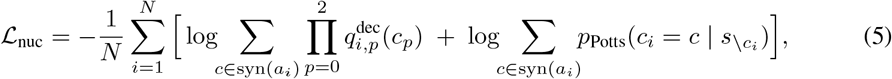

where *a*_*i*_ is the native amino acid at residue *i*. This avoids explicitly supervising arbitrary codon choices, although it does not enforce equal energies across synonymous encodings. The codon-level marginal is the three-position analogue of the site conditional (2); its normaliser sums over |Σ|^3^ = 64 codons and remains tractable. To prevent overfitting of the Potts loss at low masking rates, we apply a noise-dependent weight *ω*(*t*) = (1 *− t*)^*α*^, with hyperparameter *α >* 0, that down-weights the easy high-*t* regime, where most input residues remain visible (Appendix D.1).

### 3.3 Sampling and guidance

We use DLMC [30] to explore the discrete nucleotide space through energy-informed proposals. The chain maintains a complete sequence throughout sampling, allowing nucleotide-level objectives to be evaluated over the full sequence and combined with the structural energy. At inference time, DLMC computes per-site energy gaps Δ*E*_*µ*_(*b*) = *E*_*µ*_(*b*) *− E*_*µ*_(*s*_*µ*_) for all candidate states *b ∈* Σ, where 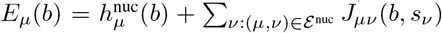 is the site-*µ* contribution to the Potts energy with the rest of *s* held fixed. These gaps construct local transition probabilities and let the chain efficiently explore the combinatorial sequence space while respecting the Potts energy landscape.

NuCaliby incorporates sequence-level objectives during sampling through proposal guidance and / or acceptance guidance. Proposal guidance uses local energy changes, or approximations to them, to favor candidate nucleotide substitutions. Acceptance guidance evaluates objectives on complete current and proposed sequences without requiring local substitution scores. The choice depends on the information available from an objective and the cost of evaluating it (Figure 2).

For proposal guidance, let *x* = onehot (*s*) and let *F* (*x*) be a continuous extension of an auxiliary sequence energy. We use its derivatives at the current sequence to augment the Potts energy gaps:

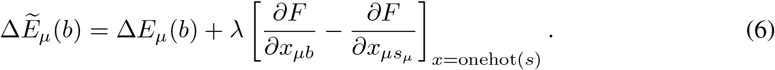

Here, *λ* controls guidance strength. The derivative difference gives a local linear estimate of the auxiliary energy change when replacing *s*_*µ*_ with *b*. For tAI and stop-codon energies, this equals the exact single-nucleotide energy change with all other positions fixed. If gradients are used, they are computed directly with respect to the one-hot representation.

We demonstrate the mechanism with the following composable energy biases:

#### tAI guidance

The tRNA Adaptation Index (tAI) [8] estimates how well a codon matches the host organism’s tRNA pool by considering the tRNA gene copies while correcting for non-standard tRNA pairing at the third codon position, called wobble correction. It is commonly used as a proxy for adaptation of coding sequences to the host tRNA pool and correlates with translational efficiency and protein expression [21, 2]. We use the length-normalized energy

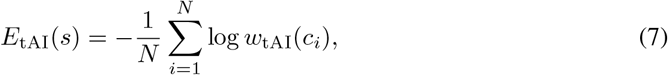

weighted by *λ* during guidance. The implementation assigns neutral log weights to start and stop codons, with stops handled by a separate penalty. Expressing each codon’s contribution as a multilinear function of its nucleotide one-hot entries allows exact single-nucleotide energy differences to be obtained through the derivative differences in Eq. 6.

#### mRNA sequence motif guidance

A compelling application of nucleotide-level design is embedding functional RNA motifs such as frame-shifting elements directly into the coding sequence. When the motif has no preferred position in the coding sequence, we use a sliding-window energy bias that lets the sampler find the best location (Figure 3).

**Figure 3.**
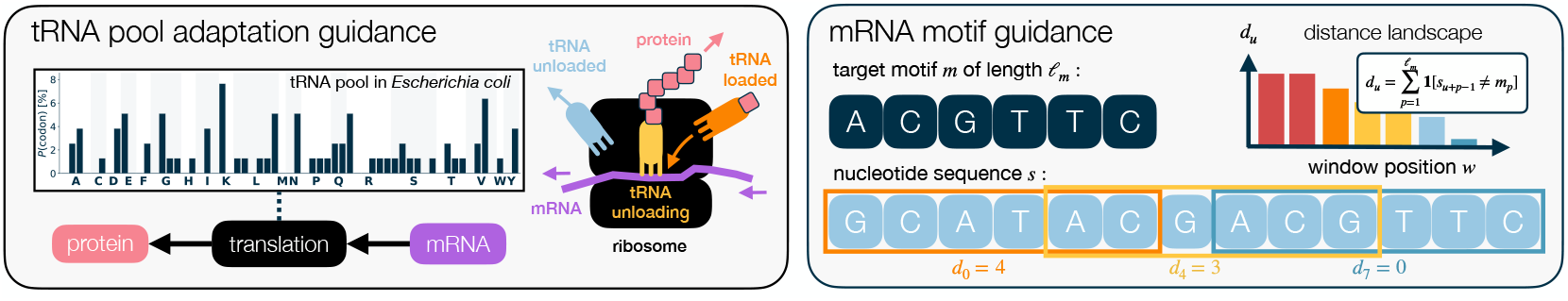
Auxiliary energy guidance terms that can improve tRNA pool adaptation for a given organism (left) or include an arbitrary mRNA motif into the coding sequence of a protein (right).

For a target motif 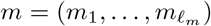 and designed sequence *s*, the Hamming distance at window position 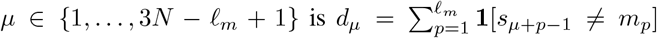 represented in one-hot coordinates as 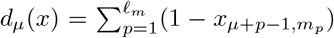. We aggregate over all windows using a smoothmin:

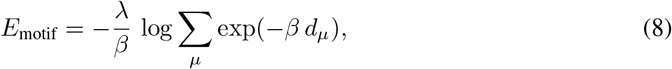

where *β* controls the sharpness (*β→ ∞* recovers the hard min over windows; smaller *β* flattens the weighting across windows) and *λ* controls the weight relative to the Potts energy. We differentiate this smooth-min energy with respect to the current one-hot sequence and use derivative differences to guide nucleotide proposals. These provide local linear approximations to changes in the motif energy. The motif energy already includes its guidance weight *λ*, which is applied only once. Multiple motifs can be placed at once by summing their individual *E*_motif_ terms, requiring no additional sampling runs.

Acceptance guidance evaluates auxiliary objectives on complete current and proposed sequences, without requiring local substitution scores. When acceptance/rejection is enabled, the proposed sequence *s*′ is accepted under the Metropolis-Hasting criterion, with probability

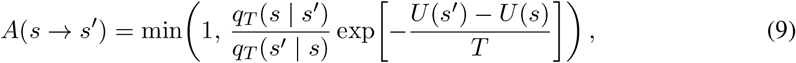

where *T* is the current sampling temperature, *q*_*T*_ is the proposal probability, and *U* includes the Potts energy and all weighted auxiliary energies used for acceptance. Auxiliary energies used to construct proposals also contribute to *U*; additional objectives can enter *U* without altering the proposal.

Our experiments use annealed DLMC without acceptance/rejection except when applying ViennaRNA acceptance guidance (Appendix A.1).

#### mRNA secondary structure guidance

We use the ViennaRNA [19] ensemble free energy per nucleotide as an acceptance-only objective. This favors sequences with more favorable predicted mRNA folding energies, while proposals remain guided by the Potts energy and stop-codon penalty. Only complete-sequence energy evaluations are required.

## 4 Experiments

Our experiments establish one claim: NuCaliby enables joint optimization across the central dogma (protein, mRNA, and DNA) with explicit control over the trade-offs between protein designability and nucleotide-level objectives. Each experiment isolates a separate facet of this claim. **Codon-level organization** (§4.1) tests whether the Potts model recovers codon structure from purely pairwise nucleotide couplings, despite operating at the level of individual nucleotides. **Designability** (§4.1) shows that nucleotide-level operation does not reduce *in silico* self-consistency of the designed proteins. **tRNA adaptation** (§4.2) illustrates a controllable trade-off between a nucleotide objective and structural compatibility, including comparison with synonymous optimization. **RNA motif inclusion** (§4.3) tests direct enforcement of sequence constraints whose compatible amino-acid patterns grow combinatorially in a motif-aware residue-level formulation. **Predicted RNA folding energy** (§4.4) demonstrates how a complete-sequence evaluator can guide acceptance without supplying local substitution scores. Finally, **wet-lab validation** (§4.5) assesses whether guided designs, including motif-containing sequences, remain expressible and soluble.

Experiments use 183 soluble monomer backbones from the Caliby test set (Appendix A.6). NuCaliby is trained on *~* 231k PDB structures from the Boltz-1 [36] training set using a temporal split, clustered at 40% sequence identity with MMseqs2 [29] and excluding transmembrane proteins. We compare against ProteinMPNN, CodonMPNN, and Caliby. Since NuCaliby learns to predict the weights of both a residue-level (AA) and a nucleotide-level (NT) Potts model, we evaluate sampling from each independently. For Caliby and NuCaliby, we additionally report ensemble-conditioned results (ENS) with Potts parameters aggregated across the original structure plus 15 variants generated by Protpardelle-1c [5] via partial diffusion. See Appendix A.1, Table 3 for guidance settings.

**Table 3:**
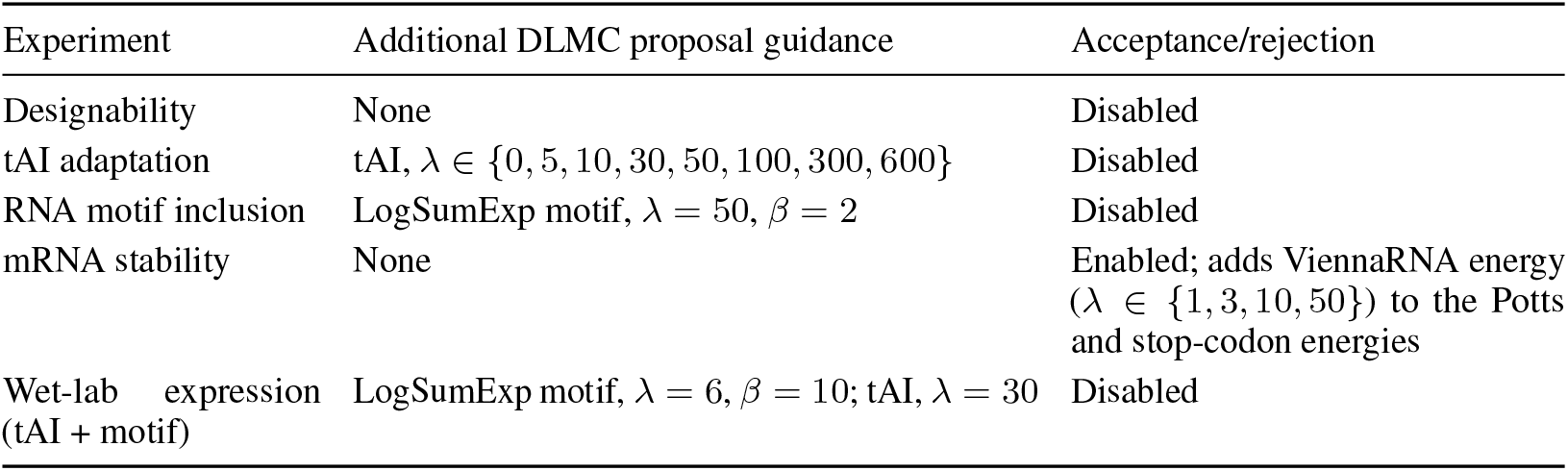
Guidance settings for NuCaliby nucleotide-level sampling. All rows share the Potts and stop-codon terms described above. The wet-lab row describes the combined tAI-plus-motif condition; the separate tAI-only conditions are detailed in Appendix B.

| Experiment | Additional DLMC proposal guidance | Acceptance/rejection |
| --- | --- | --- |
| Designability | None | Disabled |
| tAI adaptation | tAI, $\lambda \in \{0, 5, 10, 30, 50, 100, 300, 600\}$ | Disabled |
| RNA motif inclusion | LogSumExp motif, $\lambda = 50, \beta = 2$ | Disabled |
| mRNA stability | None | Enabled; adds ViennaRNA energy ($\lambda \in \{1, 3, 10, 50\}$) to the Potts and stop-codon energies |
| Wet-lab expression (tAI + motif) | LogSumExp motif, $\lambda = 6, \beta = 10$; tAI, $\lambda = 30$ | Disabled |

### 4.4 Codon-level organization and protein designability

The Potts model includes only pairwise (2-body) interactions, while codons involve constraints over nucleotide triplets. We ask whether codon structure is captured through pairwise couplings. We measure interaction strength between NT sites *µ* and *ν* as the Frobenius norm of the coupling matrix, *∥***J**_*µν*_*∥*_*F*_.

For pairs with the same sequence distance, coupling strength is higher for codon-aligned pairs than for frame-shifted pairs. This suggests that the model organizes pairwise interactions into codon-aligned triangles. Within codons, couplings between the first and second nucleotide positions are strongest, while those involving the third position are weaker, consistent with the wobble position (Figure 4).

**Figure 4.**
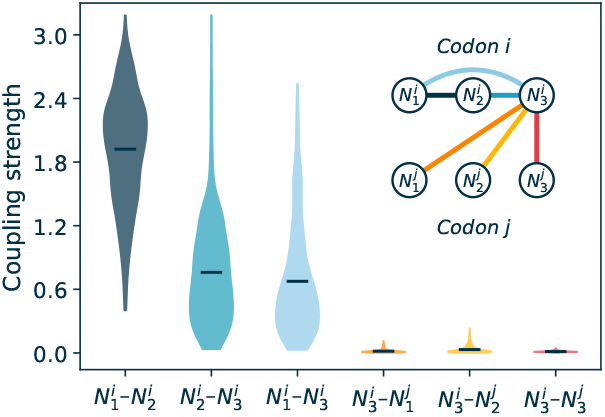
Codon-level organization in the Potts model arises from a triangle of within-codon couplings.

Designability is assessed via AlphaFold2 self-consistency (Table 1; see Appendix A for protocol). The baselines include folding with single-sequence AF2 for the (Native) sequence, ProteinMPNN, CodonMPNN and Caliby. Structure prediction for Native is challenging compared to re-designed sequences, which is a well described limitation of the current in-silico evaluation strategies [16]. Sequence recovery measures similarity to the native sequence, whereas inverse-folding design seeks sequences compatible with the target backbone. Lower recovery therefore does not by itself indicate poorer design, and we interpret it alongside structural self-consistency and experimental validation.

NuCaliby matches Caliby on sequence recovery and designability while additionally enabling nucleotide-level design. Both sampling from the residue-level Potts (NuCaliby^AA^), and the nucleotide-level Potts (NuCaliby^NT^), preserve inverse-folding performance, and ensemble conditioning further improves designability. CodonMPNN, by contrast, retains ProteinMPNN-level recovery but pays a designability cost consistent with training on predicted structures (Appendix F).

#### Ablations

We assess the contributions of nucleotide-graph connectivity and joint amino-acid/nucleotide training (Table 2). Removing either intra-codon or inter-codon edges reduces nucleotide-level designability, supporting the inclusion of both interaction types. Training the nucleotide branch alone produces a smaller decrease in performance, indicating that joint supervision benefits nucleotide-level design. Full results and experimental details are provided in Appendix E.

### 4.2 Adaptation to organism-specific tRNA pools

We evaluate sequences generated by CodonMPNN, Caliby + DNA Chisel and NuCaliby^NT^ using the same *E. coli* BL21(DE3) tRNA adaptation index (tAI) [8]. 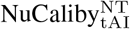 applies these tAI weights as an inference-time energy bias with strength *λ*, while DNA Chisel optimizes tAI within the synonymous encodings of Caliby-generated amino-acid sequences. CodonMPNN uses species-level *E. coli* conditioning because BL21(DE3) is sparsely represented in its training data (Appendix A.3).

Figure 5 (left) shows the trade-off between tAI and designability. Synonymous tAI optimization with Caliby + DNA Chisel achieves a mean tAI of 0.572. NuCaliby can extend this trade-off toward higher tAI under stronger guidance, reaching 0.660 at *λ* = 300 and 0.725 at *λ* = 600, at an increasing cost to self-consistency. pLDDT remains comparatively stable, while decreasing TM-scores indicate structural drift from the input backbone. Thus, synonymous optimization already provides substantial tAI gains for a fixed amino-acid sequence, while joint amino-acid and nucleotide design provides access to a broader trade-off.

**Figure 5.**
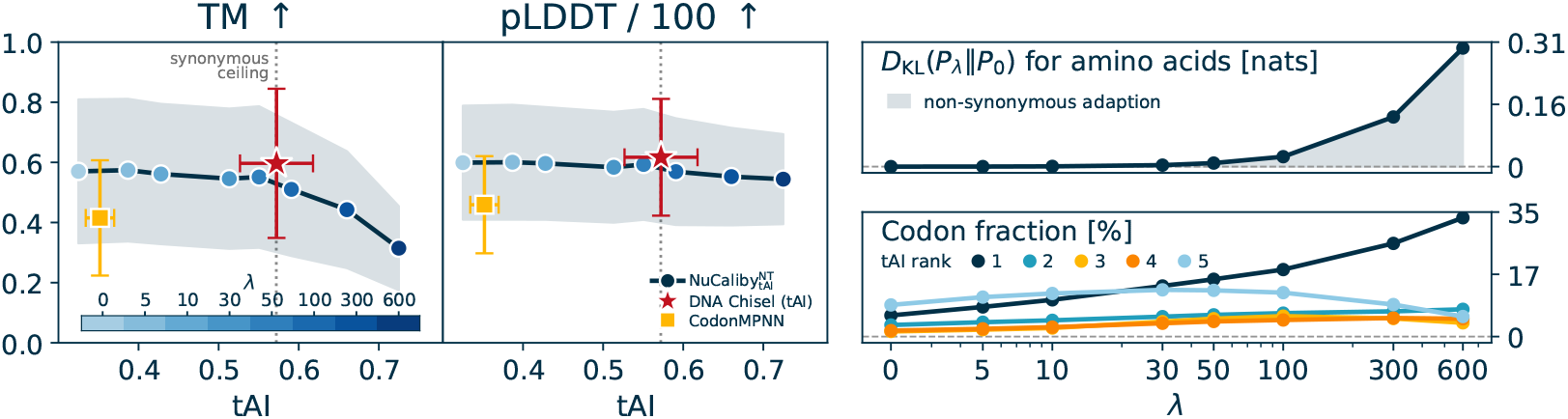
tRNA adaptation index (tAI) and in-silico designability for NuCaliby with increasing tAI guidance strength *λ*, compared with CodonMPNN and Caliby + DNA Chisel with synonymous tAI optimization. All methods are scored using the same BL21(DE3) tAI weights. Right: amino acid KL divergence relative to unbiased sampling and codon-rank frequencies as a function of *λ*.

Figure 5(right) measures the KL divergence between the amino-acid distribution under bias strength *P*_*λ*_ and the unbiased distribution *P*_0_. Up to *λ* = 30, amino-acid composition changes little, although this aggregate statistic does not establish that individual substitutions are synonymous. Stronger guidance shifts amino-acid composition and increases the frequency of rank-1 codons as self-consistency declines. These results are consistent with exploiting non-synonymous degrees of freedom to improve tAI beyond the optimized Caliby baseline. The baseline value of 0.572 is specific to those amino-acid sequences, rather than a universal limit on synonymous optimization.

### 4.3 Inclusion of mRNA sequence motif

We demonstrate motif guidance by embedding two RNA elements into designed coding sequences: SiRA aptamer [35] and HIV-1 frameshifting element (FSE) [14, 23], explained in Appendix A.4.

To construct a residue-level baseline, we translate the RNA motif into a set of candidate amino-acid patterns. The number of codon variants is bounded by *ℓ*_*m*_ mod 3 across reading frames; stop-codon-containing variants are excluded, and the remainder translate via *τ* (see problem statement in §3) to a smaller set of amino-acid sequences (Figure 10). The SiRA aptamer and HIV-1 FSE yield 264 and 129 codon variants that collapse to 68 and 31 amino-acid patterns respectively. The baseline samples sequences from Caliby and checks whether any contains one of these patterns. 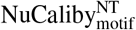 instead applies the motif energy bias (Eq. (8)) at *λ*=50, *β*=2 directly at the nucleotide level, with a stop-codon penalty (weight 100) to prevent internal stops.

Figure 6 shows that residue-level sampling does not solve the problem: none of the 183 test proteins contain the motif by chance at *n*=16. NuCaliby reaches *d*=0 on all 183 proteins by *n*=2 (single motif) and *n*=16 (two motifs); designability of motif-embedded designs improves with *n*.

**Figure 6.**
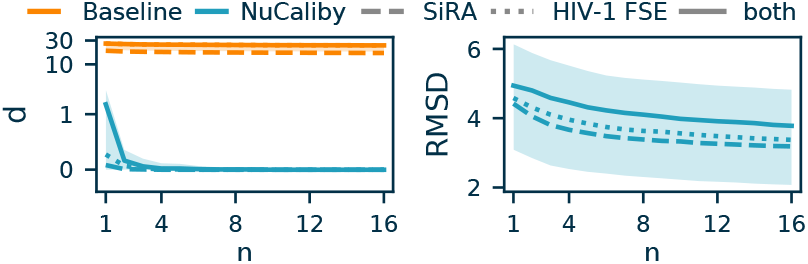
Best-of-*n* motif inclusion on 183 test monomers, for SiRA aptamer, HIV-1 FSE and both simultaneously. *d*: minimum Hamming distance to the motif. pLDDT and TM are zero when a sample has *d>*0.

Figure 7 traces how the embedded motif emerges during a single sampling run: the position of its best-matching window narrows progressively as the temperature anneals and the motif weight ramps up, with most independent trajectories converging to the same location and to an exact match. Sensitivity analyses across guidance strengths, sampling temperatures, sampling budgets, and motif choices are reported in Appendix A.5.

**Figure 7.**
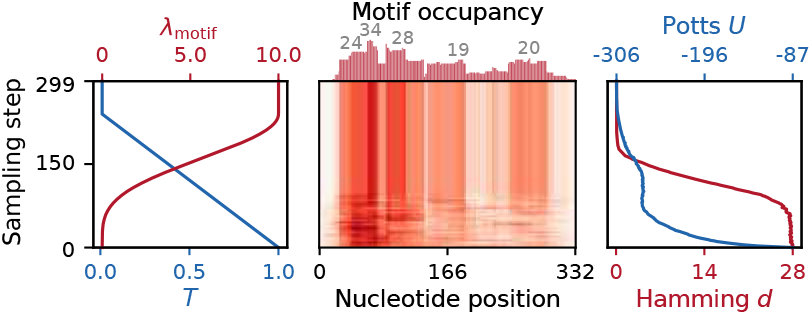
Heatmap of motif occupancy during sampling. Position of the best-matching SiRA aptamer window across 100 independent sampling trajectories where 93/100 runs converge to *d*=0.

### 4.4 mRNA folding energy optimization

We use the ViennaRNA [19] partition-function ensemble free energy per nucleotide as an acceptance-only objective (Eq. 9). As shown in Figure 8, the designs migrate toward more-negative free energy. At *λ*=10, designs for 182 of 183 backbones have lower predicted ensemble free energy per nucleotide than the unguided baseline, with a mean shift of *−*0.07 kcal*/*(mol *×* nt). This demonstrates that complete-sequence energy evaluations can guide design without supplying local substitution scores.

**Figure 8.**
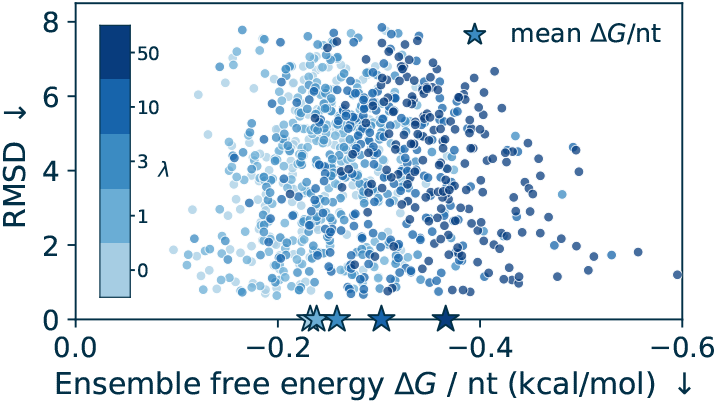
Self-consistency RMSD versus predicted mRNA ensemble free energy per nucleotide. Guidance shifts designs toward lower predicted RNA folding energies while preserving comparable self-consistency.

### 4.5 In vitro wet lab validation

We experimentally evaluated three capabilities of NuCaliby in vitro: (i) whether it can directly generate expressible DNA sequences without post-hoc codon optimization, (ii) whether inference-time tAI guidance improves protein yield, and (iii) whether RNA sequence motifs can be embedded into coding sequences while retaining expression.

Eight structurally diverse backbones were tested, with 24 sequences sampled per backbone before filtering. Unless excluded due to synthesis constraints, two designs per method and backbone were evaluated for expression and monodispersity using size-exclusion chromatography traces.

Overall, the experiments provide an initial indication that NuCaliby can generate soluble and expressible designs across unbiased, tAI-guided, and multi-objective tAI / RNA-motif-conditioned settings (Figure 9). For moderate tAI guidance *λ* = 30, soluble yield increased across all eight backbones. After averaging the two designs per backbone, the median yield increased by approximately 24%, from 0.042 to 0.052 mg/mL. This improvement was significant in an exact one-sided Wilcoxon signed-rank test (n = 8 paired backbones, W = 36, p = 0.0039). Motif-conditioned designs remained expressible in several cases, indicating that RNA-level constraints can be incorporated into coding sequences without fully abolishing protein production. CodonMPNN was overall low in yield and additionally suffered from synthesizability issues, which reduced the number of experimentally testable constructs (See Appendix B for more details).

**Figure 9.**
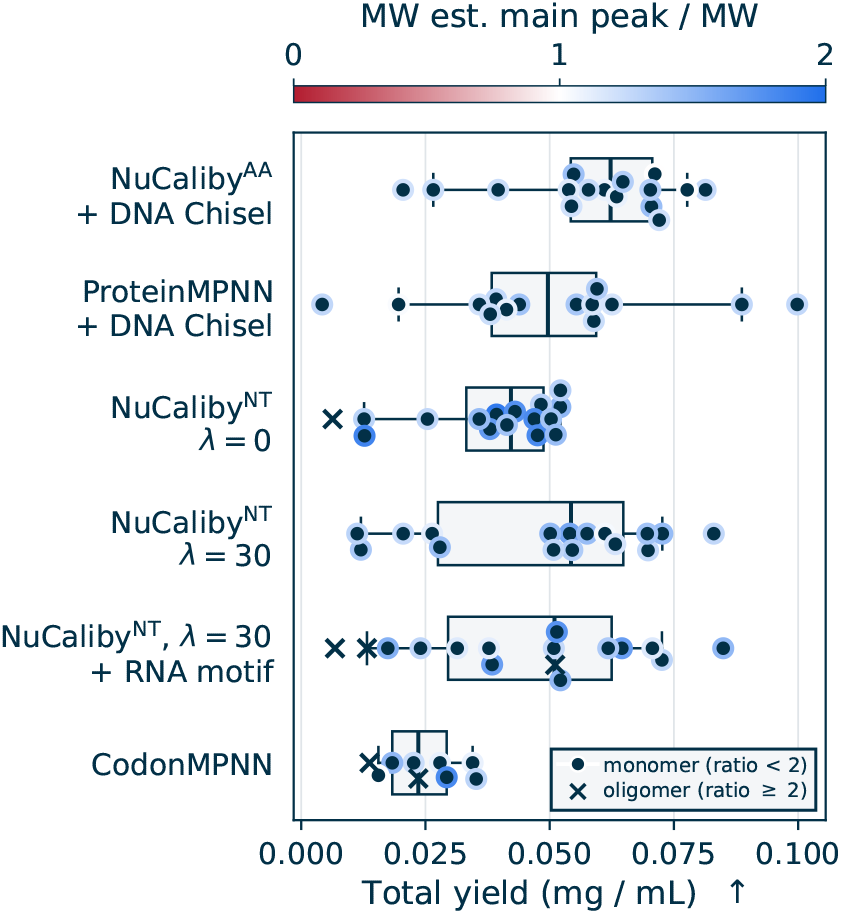
Experimental protein expression. Yields estimated from SEC traces. *λ* indicates tAI guidance strength. X indicates higher-order oligomeric state.

**Figure 10.**
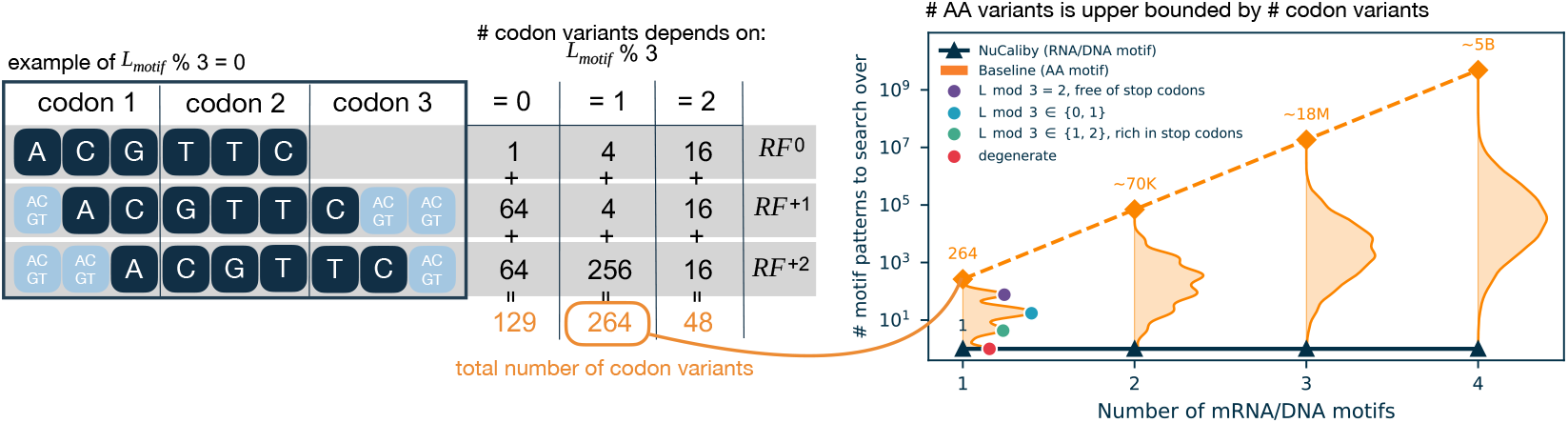
A residue-level baseline (orange) must enumerate every distinct AA pattern that the motifs produce across the three reading frames (*RF* ^0^, *RF* ^+1^, *RF* ^+2^). Theoretical worst case is 264^*k*^, attained at *ℓ*_*m*_ mod 3 = 2 free of stop codons. Densities at each number of motifs *k* summarize 1,000 random RNA motifs of length 30–100 nt (products of *k* i.i.d. samples for *k >* 1). The degenerate case means there is only one possible amino acid variant. NuCaliby adds each motif as a nucleotide-level energy term, keeping the search at a single design problem regardless of *k*.

## 5 Conclusion

NuCaliby is a structure-conditioned nucleotide-level Potts framework that jointly optimizes aminoacid identity and nucleotide coding sequence while maintaining structural consistency. Its energy-based formulation incorporates external constraints at both levels during design, allowing us to realize nucleotide objectives that are infeasible to achieve through synonymous codon choice alone.

Synonymous marginalization encourages approximate invariance to codon choice, while inference-time guidance enables explicit control over nucleotide-level objectives. Guidance can shape nucleotide proposals using local objective information or assess complete candidate sequences through acceptance, allowing multiple sequence-level objectives to be combined during design.

We show that nucleotide-level design embeds RNA sequence motifs directly into coding sequences and adapts codon usage to host tRNA pools, while retaining good self-consistency. Post-hoc codon optimization restricted to synonymous substitutions cannot satisfy these constraints: identifying potentially compatible amino-acid patterns requires combinatorial enumeration that scales poorly. ViennaRNA guidance lowers predicted mRNA ensemble free energy while preserving comparable self-consistency. Wet lab experiments on a diverse set of de novo protein backbones demonstrate that NuCaliby achieves performance on par with state of the art protein design methods like ProteinMPNN[7] producing expressible and soluble proteins, while performing a more complex optimization task.

Future directions include finer-grained guidance and a broader range of RNA elements. Natural coding sequences exhibit position-dependent behavior, such as the low-tAI ramp at the 5’ end [33]; targeting such position-specific tAI profiles, rather than globally maximizing tAI, and incorporating proxies for RNA secondary structure formation, would allow a more biologically nuanced control of translation and cotranslational protein folding [20]. We also plan to incorporate a broader range of regulatory and functional RNA elements directly into coding sequences: programmed ribosomal frameshift motifs or translational stalling signals could be embedded to modulate protein folding or generate multiple protein products from a single transcript. Although we primarily demonstrate our approach on de novo design, the same framework applies to existing proteins, with limited amino acid substitutions to satisfy nucleotide-level properties.

Nucleotide-level design combines constraints along the central dogma of molecular biology. As a proof-of-concept, NuCaliby points toward protein design as a multi-scale optimization problem spanning amino-acid structure, nucleotide sequence, and cellular context.

## Acknowledgments

The authors thank Michael A. Jendrusch for helpful discussions, and Nele Dierlamm and Emma Weißert for help with experiments. L.J.D. and L.F.M. are grateful for funding by the Deutsche Forschungsgemeinschaft (DFG) – Project number 529882161. This work was funded under Germany’s Excellence Strategy – EXC3092 – 533751719.

## A. Experimental details

NuCaliby experiments use a single checkpoint trained on the Boltz-1 training split (see Appendix D.1 for training details). Sampling is performed with DLMC for 500 sweeps with the Potts temperature annealed to a final value *T*_final_=0.01. Designability is evaluated by folding the designed amino-acid sequence with AlphaFold2 [15] via ColabFold [22] in single-sequence mode with 4 recycles using model 1, and comparing predicted structures to the input backbone with TM-align [41].

### Interpreting sequence recovery

Native-sequence recovery is a useful reference metric, but the native sequence is not the only possible solution to an inverse-folding task. Furthermore, high sequence recovery does not relate to high self-consistency [3].We therefore report sequence recovery alongside self-consistency and prediction confidence, and use expression and solubility measurements to assess experimental feasibility. Neither high nor low recovery alone establishes design quality.

### A.1 Guidance settings

Table 3 summarizes the guidance used in the main nucleotide-level design experiments. In every listed setting, the DLMC proposal includes the nucleotide Potts energy with weight 1 and a premature stop-codon penalty with weight 100. The table lists additional proposal terms and indicates whether Metropolis–Hastings (M–H) acceptance/rejection is enabled. “None” denotes no additional proposal guidance.

When enabled, acceptance uses the total energy, including terms that also influence proposal construction. The *λ* = 0 RNA comparison uses the unguided baseline.

### A.2 Compute resources

Training data preprocessing with AtomWorks [27] takes 10–20 hours on a multi-CPU node. Pre-training NuCaliby end-to-end takes approximately 15 hours on 8 *×* NVIDIA B200 GPUs. At inference, the encoder predicts the Potts parameters for a single backbone in well under one second on a single H100 GPU; subsequent DLMC sampling reaches roughly 10^3^ sweeps per second for proteins of *~*240 residues, so 500 sweeps complete in *<* 1 s and a single design takes *~*1–2 s end-to-end. Sampling the full 183-protein test set therefore takes only a few minutes per configuration. Structure prediction with ColabFold AF2 in single-sequence mode runs on commodity CPU servers; a GPU is not required, and parallelising independent designs across CPU workers makes evaluation throughput essentially proportional to the available core count.

### A.3 Adaptation to organism-specific tRNA pools

The tAI weights for *E. coli* BL21(DE3) (NCBI tax ID 469008) are computed from the host’s tRNA gene copy numbers following [8]. We sweep the bias strength *λ ∈ {*0, 5, 10, 30, 50, 100, 300, 600*}* and sample one sequence per protein at each setting. All methods are evaluated on the same 183 test backbones using a best-of-1 sampling budget and the same BL21(DE3) tAI weights. NuCaliby uses these weights for inference-time guidance, and DNA Chisel uses them to optimize synonymous encodings of Caliby-generated amino-acid sequences for tAI. CodonMPNN is conditioned on the species-level *E. coli* label (NCBI tax ID 562), represented by approximately 21,000 training examples, rather than BL21(DE3), represented by only one example. The tRNA gene copy numbers used to compute the tAI weights are strain-specific. Thus, the evaluation objective is shared, while organism conditioning follows the information available to each method.

### A.3 mRNA sequence motif inclusion

#### Motifs

**SiRA aptamer [35]** is a synthetic RNA aptamer that binds silicon-rhodamine fluorophores and has previously been inserted into untranslated regions for live-cell RNA imaging using super-resolution microscopy. Here, it serves as a representative example of a complex RNA sequence motif. We do not evaluate the downstream functional consequences of motif embedding on translation efficiency **HIV-1 frameshifting element (FSE) [14, 23]** is a short RNA element embedded within the HIV coding sequence that induces programmed ribosomal −1 frameshifting. This causes the ribosome to shift into an alternative reading frame, enabling a single mRNA to encode multiple protein products.

#### NuCaliby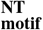

We sample with the motif energy bias (Eq. (8)) at *λ*=50 and *β*=2, and additionally apply a stop-codon penalty with weight 100 to keep the embedded motif from introducing internal stops. The two-motif setting sums the per-motif energies. We sample *n*=16 sequences per protein and report best-of-*n* success.

Motif placement is robust across a broad range of (*λ, β*) values: in our experiments the model achieves a perfect single-motif embedding (*d*=0) within at most 2 samples and a two-motif embedding within at most 16 samples on every tested backbone. We therefore use slightly different settings in different experiments (e.g. *λ*=6, *β*=10 in the wet-lab pipeline of Appendix B, where we additionally combine motif guidance with *λ*_tAI_=30) without observing material differences in success rate.

#### Caliby AA-level baseline

We sample *n*=64 residue-level sequences per protein from Caliby and check whether any translation matches one of the amino-acid variants derived from the motif across the three reading frames (Figure 10).

### A.5 Sensitivity to sampling hyperparameters

We assess motif inclusion on 183 test monomers using one sample per configuration (Table 4). Each configuration varies one setting from the reference configuration: SiRA motif, guidance strength *λ* = 50, smooth-min sharpness *β* = 2, initial temperature *T*_init_ = 1.0, final temperature *T*_final_ = 0.01, and 500 DLMC sweeps. Temperature is annealed over the first 80% of sweeps. We report the percentage of designs with an exact motif match (*d* = 0) or a match within one nucleotide (*d ≤* 1), together with TM and pLDDT.

**Table 4:** Sensitivity of motif inclusion to sampling hyperparameters and motif identity. Each row uses one sample per test backbone. The two inclusion columns are percentages; scTM and pLDDT are mean designability metrics. Settings other than the one indicated follow the reference configuration described above.

| Parameter | Setting | $d = 0$ (%) ↑ | $d \leq 1$ (%) ↑ | TM ↑ | pLDDT ↑ |
| --- | --- | --- | --- | --- | --- |
| Guidance strength and smooth-min sharpness |  |  |  |  |  |
| $\lambda$ | 25 | 87.4 | 100.0 | .487 | 52.6 |
| $\lambda$ | 50 | 91.8 | 100.0 | .503 | 53.4 |
| $\lambda$ | 100 | 87.4 | 100.0 | .490 | 53.0 |
| $\beta$ | 1 | 90.2 | 100.0 | .484 | 52.7 |
| $\beta$ | 2 | 91.8 | 100.0 | .503 | 53.4 |
| $\beta$ | 5 | 92.3 | 100.0 | .500 | 53.6 |
| Sampling temperature |  |  |  |  |  |
| $T_{\text{final}}$ | 0.005 | 92.3 | 100.0 | .494 | 53.0 |
| $T_{\text{final}}$ | 0.01 | 91.8 | 100.0 | .503 | 53.4 |
| $T_{\text{final}}$ | 0.05 | 91.3 | 100.0 | .492 | 53.2 |
| $T_{\text{init}}$ | 0.5 | 94.5 | 100.0 | .491 | 53.0 |
| $T_{\text{init}}$ | 1.0 | 91.8 | 100.0 | .503 | 53.4 |
| $T_{\text{init}}$ | 2.0 | 74.9 | 100.0 | .496 | 53.3 |
| Sampling budget |  |  |  |  |  |
| DLMC sweeps | 250 | 87.4 | 100.0 | .493 | 53.7 |
| DLMC sweeps | 500 | 91.8 | 100.0 | .503 | 53.4 |
| DLMC sweeps | 750 | 89.1 | 100.0 | .495 | 53.4 |
| Motif identity and length |  |  |  |  |  |
| Motif | SiRA (50 nt) | 91.8 | 100.0 | .504 | 53.4 |
| Motif | FSE (66 nt) | 72.7 | 100.0 | .465 | 50.3 |
| Motifs | SiRA + FSE (50 + 66 nt) | 61.2 | 88.0 | .410 | 47.4 |

Within the tested ranges, guidance strength, smooth-min sharpness, final temperature, and sampling budget have relatively small effects on designability, and all designs remain within one nucleotide of the target motif. Initial temperature has a larger effect on exact inclusion: increasing *T*_init_ from 1.0 to 2.0 lowers the exact-match rate from 91.8% to 74.9%. Motif choice also affects inclusion and designability. Simultaneous SiRA and FSE inclusion achieves 61.2% exact matches in best-of-1 sampling, with lower TM and pLDDT than either motif alone, although all backbones achieve exact inclusion with up to 16 samples in the best-of-*n* experiment (Figure 6). Because these motifs differ in sequence as well as length, this comparison does not isolate the effect of motif length.

### A.6 List of pdbs in test set

The 183 soluble monomers from the Caliby benchmark used in all evaluations.

<preformat>

7qnl, 7qpk, 7qpl, 7qqd, 7qsj, 7sc4, 7t4w, 7tl6, 7u08, 7u71, 7uba, 7ui4, 7uk8, 7uqu, 7ur2, 7urp, 7v34, 7wkp, 7wmy, 7wnh, 7woi, 7wrn, 7x2x, 7xcc, 7xh4, 7xhz, 7xrx, 7xs3, 7xvi, 7xwy, 7xz3, 7y01, 7y1s, 7y3z, 7y43, 7y4h, 7y7o, 7y8h, 7yia, 7ykm, 7yl4, 7ynx, 7ypq, 7ysi, 7yuk, 7yyg, 7z3i, 7z64, 7zao, 7zcb, 8a0a, 8a16, 8acc, 8acr, 8adc, 8add, 8agg, 8agr, 8ahd, 8an0, 8au6, 8axj, 8ay2, 8b2e, 8b2h, 8b2s, 8b4c, 8b5w, 8b6a, 8b6p, 8b6q, 8b6u, 8bat, 8bbr, 8bbz, 8bdq, 8bf8, 8bfy, 8bl6, 8bux, 8bvi, 8bw6, 8c27, 8c4d, 8cih, 8ck0, 8cmq, 8cpk, 8cpn, 8cpq, 8cr7, 8d40, 8daj, 8deo, 8dhj, 8dvq, 8dwj, 8dx0, 8e5d, 8e5e, 8ec3, 8eel, 8efz, 8ehn, 8eil, 8eov, 8erm, 8ex9, 8exa, 8ezw, 8f5g, 8f9y, 8fkl, 8g0n, 8g32, 8g53, 8g8e, 8ga6, 8gld, 8gmx, 8gmz, 8gsx, 8gsy, 8gty, 8gxr, 8gy4, 8gyr, 8h0l, 8h18, 8h1j, 8h55, 8h5s, 8hbb, 8hbr, 8he2, 8hm4, 8hp8, 8htx, 8huz, 8i3j, 8iib, 8in6, 8iom, 8iqc, 8irq, 8j69, 8jah, 8jdg, 8jw3, 8jws, 8k5u, 8oei, 8ok3, 8olj, 8ou1, 8ow8, 8oyy, 8p0c, 8p5q, 8paq, 8pmu, 8pof, 8pxc, 8qcw, 8r8a, 8sot, 8ssu, 8sxs, 8t1m, 8t3j, 8t4c, 8t8l, 8tfs, 8tjg, 8tnm, 8tvl, 8uk7, 8upi, 8urn, 8uz8, 8v1k, 8w7p, 8y4u

</preformat>

## B Wet-lab validation

To obtain a diverse set of protein backbones for sequence generation, we curated a set of eight structures spanning different sources and design methods (Table 5). The set includes four published de novo backbones retrieved from the protein design archive [4], and four additional de novo backbones generated with RF diffusion [34]. This selection covers a range of backbone generation approaches, including AlphaFold hallucination, RFjoint2, Chroma, and RF diffusion.

**Table 5:** Protein backbones used for experimental evaluation. SSE = secondary structure estimation.

| Name | PDB | BB method | Seq method | Category | Publication | SSE | Length [#AA] |
| --- | --- | --- | --- | --- | --- | --- | --- |
| halc1_878 | 8CYK | AF hallucination | ProteinMPNN | Published de novo | Science (2022) | $\alpha/\beta$ | 136 |
| BBn6 | 9XZT | RFjoint2 | ProteinMPNN | Published de novo | PNAS (2024) | $\beta$ | 95 |
| Unc_239 | 8TNO | Chroma | Chroma | Published de novo | Nature (2023) | $\alpha$ | 285 |
| tbf_24 | 8OYS | AF hallucination | ProteinMPNN | Published de novo | Nature (2024) | $\alpha/\beta$ | 192 |
| RF1_14 | – | RF diffusion 1 | – | De novo (this work) | – | $\alpha/\beta$ | 120 |
| RF1_3 | – | RF diffusion 1 | – | De novo (this work) | – | $\alpha/\beta$ | 120 |
| RF1_2 | – | RF diffusion 1 | – | De novo (this work) | – | $\alpha$ | 120 |
| RF1_6 | – | RF diffusion 1 | – | De novo (this work) | – | $\alpha/\beta$ | 120 |

### B.1 Creating sequences

We generated 24 sequences per backbone for each method, providing an equal sampling budget across sequence design approaches:

- **ProteinMPNN** with soluble weights and default inference settings [11].
- **CodonMPNN** with default inference settings, conditioned on *E. coli* (Taxon ID 562), which constitutes 22% of the taxon IDs in its training set.
- **NuCaliby** with tAI energy biases *λ ∈ {*0, 30, 50, 300*}*.
- **NuCaliby** tAI energy bias *λ* = 30 and RNA motif guidance, embedding a random 29-nucleotide sequence: GTCCCCGCAGGTTGATCCTAAACGGCTCT.
- **NuCaliby** (amino acid branch), sampling from the residue-level Potts model.

For methods that generate amino acid sequences, namely ProteinMPNN and the NuCaliby amino acid branch, we obtained DNA sequences by reverse translation followed by codon optimization with DNA Chisel. Optimization targeted *E. coli* codon usage using CodonOptimize and MaximizeCAI, while enforcing translation, removing BsaI sites, constraining GC content to 30–70%, and requiring 80-mer uniqueness.

We applied a multi-stage filtering pipeline to ensure constructability and structural quality. First, we removed sequences containing the BsaI restriction site required for Golden Gate cloning, as well as sequences with premature stop codons. For motif-guided designs, we used a tAI guidance strength of *λ* = 30 and a motif guidance strength of *λ* = 6 with a smooth min sharpness of *β* = 10.0 and retained only sequences containing the target motif.

We then evaluated all translated amino acid sequences with AlphaFold2, 4 recycles, model 1, single-sequence mode. Designs were screened by pLDDT *>* 80 and self-consistency RMSD (scRMSD) *<* 2 Å. Filtering was applied independently for each backbone and design method. When more than three candidates passed these structural filters, we preferred designs with isoelectric point pI *<* 6.5; if this yielded fewer than three candidates, we relaxed the preference to pI *<* 6.5 or pI *>* 8.5. When fewer than three candidates passed the structural filters, we retained the top three candidates ranked by increasing scRMSD, using pLDDT as a secondary criterion.

All surviving candidates were screened using the GenScript sequence complexity check. If more than two designs remained after filtering, we selected the two candidates with the lowest self-consistency RMSD (scRMSD) for synthesis. Designs failing the GenScript screen were instead submitted to Twist fragment synthesis. Sequences that also failed Twist synthesis constraints were classified as unsynthesizable.

This procedure evaluates all methods under an equal sampling budget and a shared selection pipeline, comparing them based on their best achievable designs.

### B.2 Expression

Experimental validation procedures followed the SAPP pipeline [25] Final evaluations of protein expressibility were based on size exclusion chromatography after Ni-NTA based HIS-tag purification of a 4 mL autoinduction culture grown for 22h at 37 C. Protein yield and expected monodispersity were derived from the elution profile on a Cytiva Superdex 75 Increase 5/150 GL column in PBS at room temperature.

### B.3 Details for in-silico filtering and tested designs

#### Prefiltering sequence motifs

We started with 8 backbones *×*24 sequences = 192 sequences per method. Overall, there were 9 methods, as described in B1. Thus, 1728 sequences were subject to filtering.

Sequences that were guided to contain the motif were filtered to check whether they contained the motif:

- Motif GTCCCCGCAGGTTGATCCTAAACGGCTCT occurs in 123 out of 192 sequences (64.1%).

Further filtering removed Type IIS restriction sites reserved for the cloning reaction (commonly removed):

- GGTCTC and GAGACC.
- This removed 116 designs: 95 for CodonMPNN, 14 for NuCaliby NT, 5 for NuCaliby NT 30 tAI, and 2 for NuCaliby NT 30 tAI + Motif.

#### Filtering amino acid properties

Stronger guidance usually decreases the pass rate at the amino acid level as more constraints have to be satisfied, which is in line with the previously reported trade-off in tAI and motif inclusion. Without constraints, the pass rate of the amino acid and nucleotide level NuCaliby model is similar. We provide the filtering results below, aggregated over all backbones:

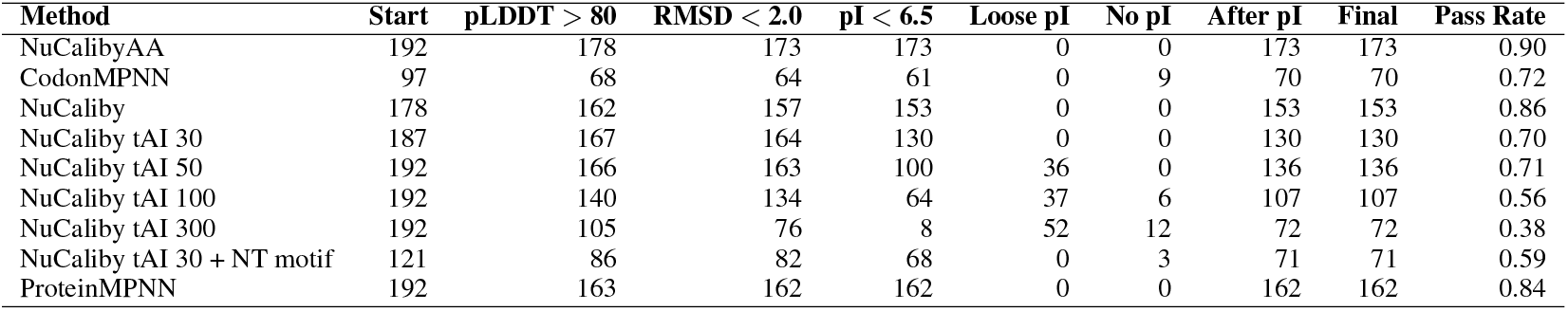

#### Filtering synthesis constraints

For each of these DNA sequences, we took all sequences that passed the filters and checked whether they could be synthesized by a commonly used gene synthesis company (GenScript). If more than two sequences per method passed, we chose those with the smallest scRMSD.

Overall, strongly guided sequences as well as CodonMPNN sequences had repetitive patterns, which made more than 50% of them too complex to be synthesized (we note that this issue did not occur for NuCaliby base). The pass rate decreased overall with increasing guidance.

| Method | Passed | Failed | Total | Pass Rate (%) |
| --- | --- | --- | --- | --- |
| NuCalibyAA | 173 | 0 | 173 | 100.0 |
| CodonMPNN | 33 | 37 | 70 | 47.1 |
| NuCaliby | 153 | 0 | 153 | 100.0 |
| NuCaliby tAI 30 | 87 | 43 | 130 | 66.9 |
| NuCaliby tAI 50 | 72 | 64 | 136 | 52.9 |
| NuCaliby tAI 100 | 47 | 60 | 107 | 43.9 |
| NuCaliby tAI 300 | 10 | 62 | 72 | 13.9 |
| NuCaliby tAI 30 + NT motif | 54 | 17 | 71 | 76.1 |
| ProteinMPNN | 122 | 40 | 162 | 75.3 |

If fewer than two sequences passed the GenScript complexity filters per backbone and method, we ordered the two sequences with the best RMSD from another gene synthesis company specializing in more complex sequences (Twist). This worked for 12 out of 29 missing designs after two synthesis attempts.

**Figure 11.**
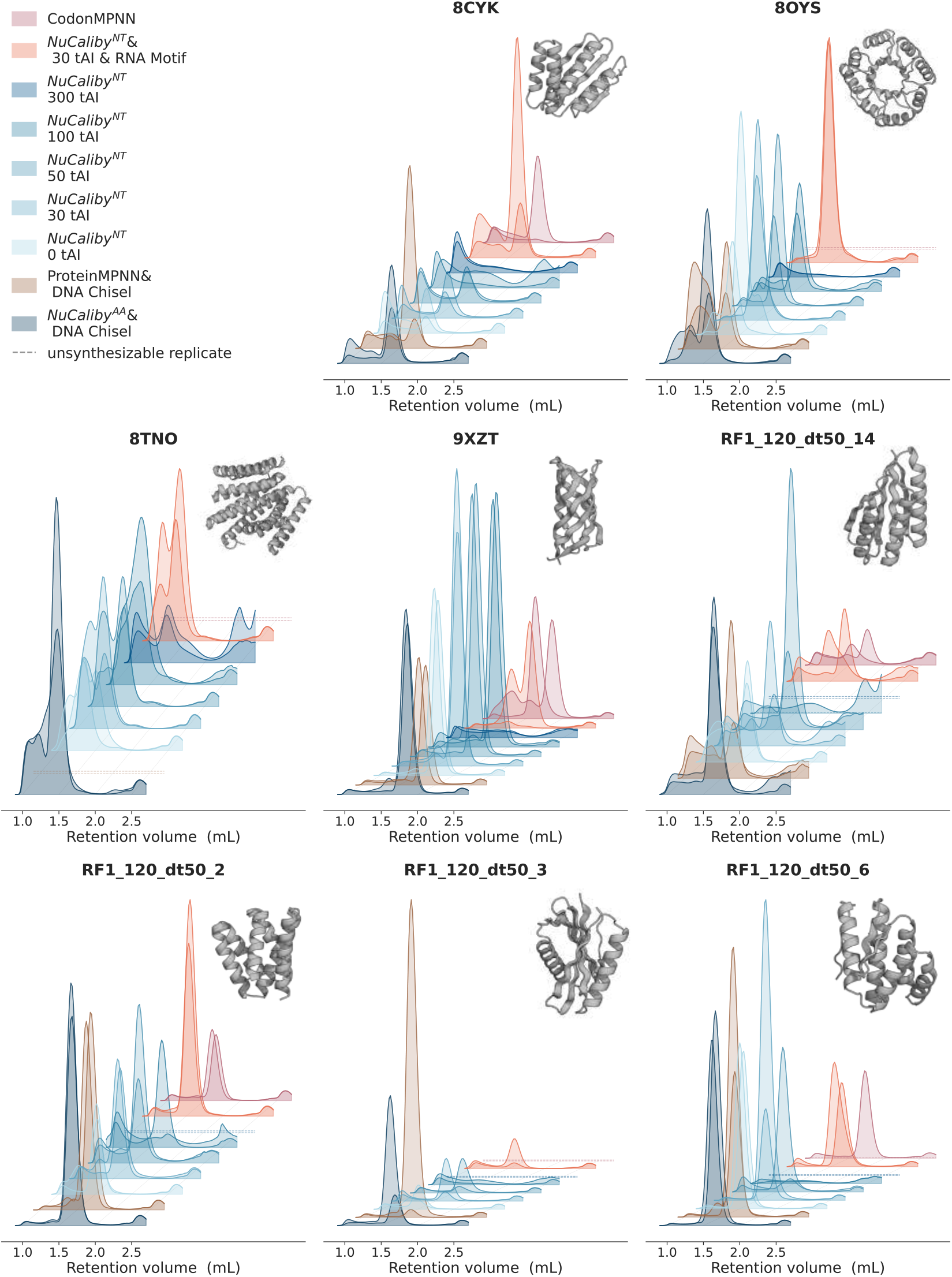
Size-exclusion chromatograms for different backbones and sequence design methods. Dashed traces indicate unsynthesizable designs for which the DNA sequence could not be synthesized as a fragment by GenScript or Twist.

Thus, overall, we obtained:

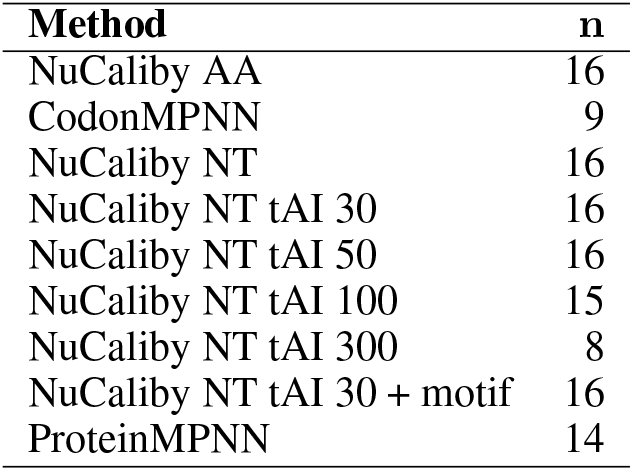

| **Method** | **n** |
| --- | --- |
| NuCaliby AA | 16 |
| CodonMPNN | 9 |
| NuCaliby NT | 16 |
| NuCaliby NT tAI 30 | 16 |
| NuCaliby NT tAI 50 | 16 |
| NuCaliby NT tAI 100 | 15 |
| NuCaliby NT tAI 300 | 8 |
| NuCaliby NT tAI 30 + motif | 16 |
| ProteinMPNN | 14 |

## C Limitations

### Approximate synonymous invariance

Ideally, the structural energy would assign equal energies to synonymous encodings of the same amino-acid sequence. We assess this property across all 183 test monomers by sampling one unguided design per backbone and evaluating 100 random single-codon substitutions of each type. The median per-protein mean absolute energy change is 0.085 [IQR: 0.073–0.095] for synonymous substitutions and 3.54 [3.33–3.67] for non-synonymous substitutions (Figure 12A). The median within-protein ratio is 2.4%, indicating substantially weaker sensitivity to synonymous changes, but not exact invariance.

**Figure 12.**
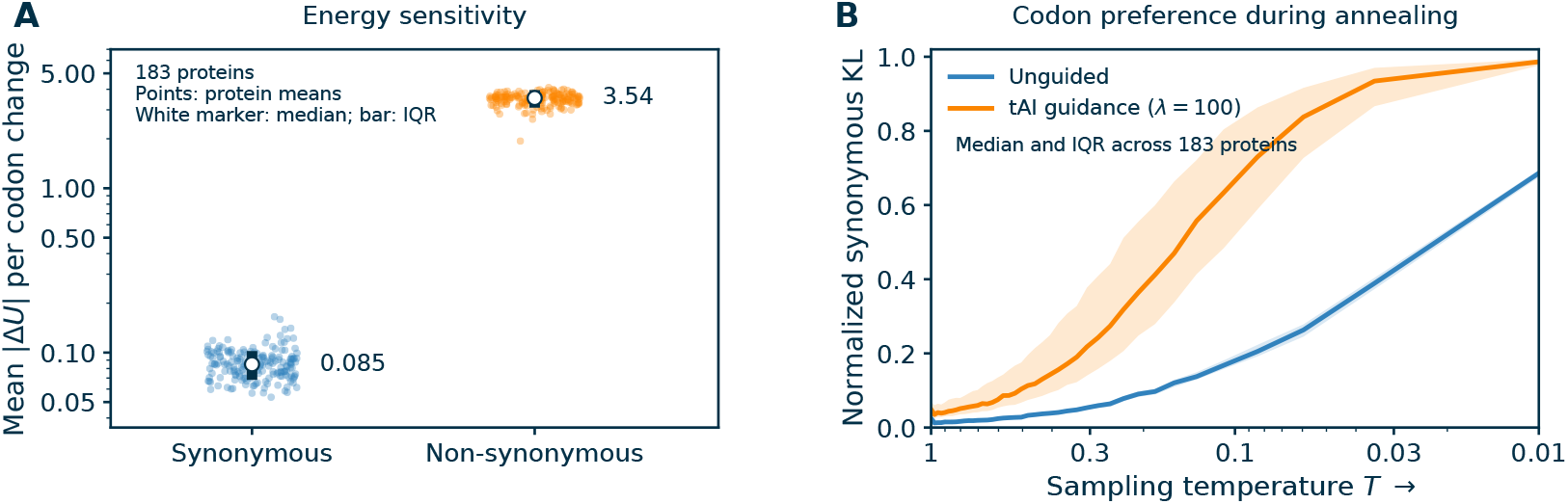
Approximate synonymous invariance across 183 test proteins. **A:** Each point is one protein’s mean absolute Potts-energy change over 100 random single-codon substitutions, shown separately for synonymous and non-synonymous substitutions on a logarithmic energy axis. White markers and dark bars indicate the median and interquartile range across proteins. **B:** Normalized KL divergence of synonymous-codon conditional distributions from uniform during unguided and tAI-guided (*λ* = 100) DLMC sampling. Lines and shaded bands show the median and interquartile range across proteins; zero denotes uniform conditionals and one denotes concentration on a single codon. Measurements are averaged over eligible residues within each protein. Repeated measurements at the final temperature are averaged within each protein before aggregation.

To assess the effect of residual preferences during sampling, we evaluate conditional distributions over synonymous codons along one unguided and one tAI-guided DLMC trajectory per backbone, each comprising 500 sweeps with temperature annealed from 1.0 to 0.01 over the first 80% of sweeps.

Every 10 sweeps, we hold the remaining sequence fixed and evaluate *p*(*c*) *∝* exp(*−U*_*c*_*/T*) over the synonymous codons of each eligible residue, including the tAI energy in the guided condition. We normalize the KL divergence from the uniform distribution by the logarithm of the synonymous-set size and average over residues with more than one synonymous codon. These preferences are weak at high temperature but become pronounced during annealing (Figure 12B): at *T* = 0.01, the median normalized divergence is approximately 0.69 without guidance and 0.99 with tAI guidance at *λ* = 100. Thus, small residual energy differences can substantially bias low-temperature codon choice. Synonymous marginalization encourages approximate invariance but does not guarantee uniform sampling over synonymous encodings.

#### Simplified biological proxies

Our nucleotide-level objectives measure tRNA adaptation, motif-sequence inclusion, and predicted RNA folding energy. The wet-lab experiments establish expression and solubility of guided designs, but do not directly measure translation efficiency, RNA motif activity, or cellular mRNA stability. Translation efficiency is governed by multiple interacting factors, including codon usage, mRNA secondary structure, and ribosome dynamics [17, 39]. Similarly, functional RNA motifs are not defined solely by their primary sequence but often depend on higher-order structural features, which are not captured by sequence-level motif incorporation alone [35]. The framework, however, is agnostic to the form of the bias term: more biologically faithful objectives (e.g. position-dependent tAI profiles, RNA secondary-structure constraints) can be plugged in via the same DLMC-with-guidance and Metropolis–Hastings interfaces without retraining.

## D Implementation details

### D.1 Noise-dependent Potts loss weighting

The Potts conditional is evaluated using the ground-truth sequence at non-target positions, while the predicted Potts parameters depend on the masked sequence supplied to the encoder. We use *t* for the probability of retaining a residue in that input, so the masking probability is 1 *− t*. At high *t*, the encoder sees nearly the full native sequence. Training and validation losses stratified by *t* show that overfitting is concentrated in this mostly unmasked regime (Figure 13). We therefore apply the weight *ω*(*t*) = (1 *− t*)^*α*^ to the Potts loss, reducing the contribution of these easy examples. This suppresses the observed overfitting and improves validation Potts loss across conditioning levels. We use *α* = 1.25.

**Figure 13.**
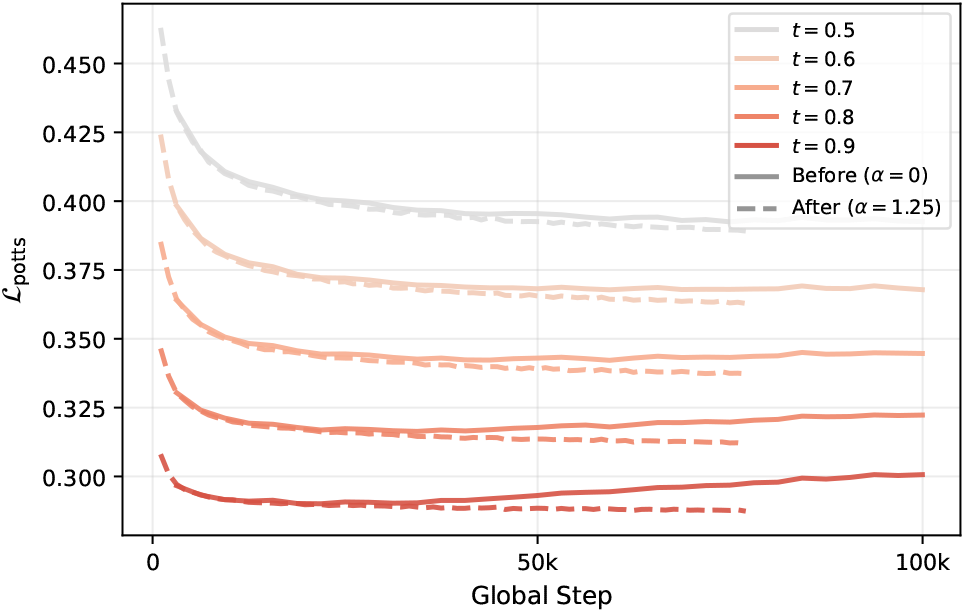
Noise-dependent Potts loss weighting helps prevent overfitting on highly unmasked sequences. Validation Potts loss stratified by sequence-retention probability *t* (masking probability 1 *− t*). Without weighting (solid), high-*t* bins overfit after *~*40k steps. With (1 *− t*)^*α*^ weighting (dashed, *α*=1.25), overfitting is eliminated and the loss improves across all *t*.

## E Ablation studies

We evaluate nucleotide-graph connectivity and joint amino-acid/nucleotide supervision on the 183 test monomers using the designability protocol in Appendix A. Tables 6 and 7 report means and standard deviations across proteins.

**Table 6:** Effect of nucleotide-graph connectivity on nucleotide-level sampling.

| NT graph | AA recovery (%) | pLDDT | scTM | scRMSD (Å) |
| --- | --- | --- | --- | --- |
| Intra + inter | $38.6 \pm 7.1$ | $60.8 \pm 19.1$ | $.579 \pm .245$ | $3.85 \pm 1.82$ |
| No intra-codon edges | $37.9 \pm 7.3$ | $55.1 \pm 18.2$ | $.504 \pm .225$ | $4.40 \pm 1.72$ |
| No inter-codon edges | $36.8 \pm 6.6$ | $54.1 \pm 17.7$ | $.500 \pm .223$ | $4.49 \pm 1.80$ |

**Table 7:** Effect of joint and separate branch training, evaluated by sampling from the indicated Potts model.

| Training | Sampling | AA recovery (%) | pLDDT | scTM | scRMSD (Å) |
| --- | --- | --- | --- | --- | --- |
| AA + NT | AA | $39.6 \pm 7.2$ | $61.8 \pm 19.6$ | $.593 \pm .246$ | $3.84 \pm 1.85$ |
| AA only | AA | $40.0 \pm 7.2$ | $62.1 \pm 19.5$ | $.603 \pm .245$ | $3.71 \pm 1.86$ |
| AA + NT | NT | $38.6 \pm 7.1$ | $60.8 \pm 19.1$ | $.579 \pm .245$ | $3.85 \pm 1.82$ |
| NT only | NT | $38.7 \pm 7.2$ | $58.9 \pm 18.5$ | $.548 \pm .227$ | $4.15 \pm 1.73$ |

### Nucleotide-graph connectivity

We train two additional models from scratch, removing either intra-codon or inter-codon edges. Without intra-codon edges, interactions among nucleotides within a codon must be mediated indirectly through neighbouring codons. Without inter-codon edges, the nucleotide Potts coupling graph separates into codon-level components; the predicted fields can still carry contextual information, but direct pairwise couplings between codons are absent. Removing either edge type reduces designability (Table 6).

### Joint AA/NT training

We compare joint training of both branches with training either branch alone (Table 7). The amino-acid branch performs similarly under joint and AA-only training. The nucleotide branch benefits from joint supervision, with higher pLDDT and scTM and lower scRMSD than NT-only training.

## F Effect of training data on designability

CodonMPNN [28] achieves comparable sequence recovery to ProteinMPNN (44.5% vs 44.4%) yet substantially lower designability (TM .441 vs .509, pLDDT 47.0 vs 54.3). CodonMPNN is trained on genomic sequences paired with AlphaFold-predicted structures rather than experimentally determined PDB structures. Tan et al. [31] show that training inverse folding models on AFDB structures degrades sequence recovery by up to 27 percentage points relative to PDB-trained models, and that even debiasing predicted structures does not fully close this gap. Our results suggest that the penalty manifests not in recovery rates but in designability, where the structural fidelity of the training data directly impacts whether designed sequences fold correctly and are self-consistent with the original backbone (Figure 14). PDB alone does not provide enough examples for different taxons to train an organism-conditioned model like CodonMPNN, motivating NuCaliby’s approach of training on PDB and deferring organism-specific preferences to inference-time energy biases.

**Figure 14.**
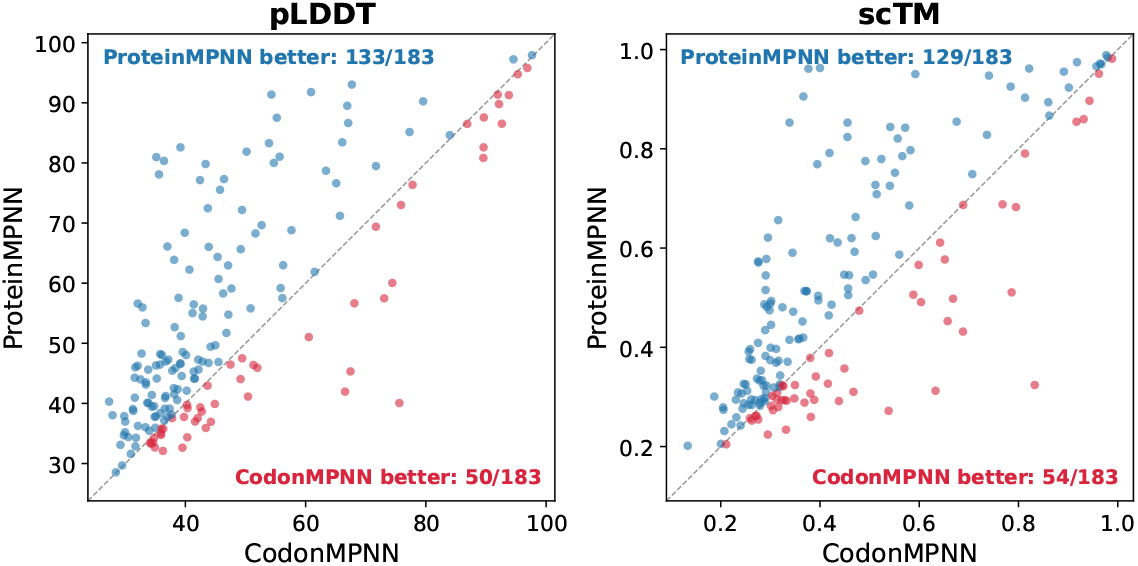
Per-protein designability comparison between ProteinMPNN trained on experimentally determined structures deposited at PDB vs CodonMPNN trained on predicted strcutures from AlphaFold Database. Each point is one of the 183 test proteins. Points above the diagonal indicate higher designability for ProteinMPNN. ProteinMPNN produces more designable sequences on 133/183 proteins by pLDDT and 129/183 by scTM, suggesting that training on predicted structures degrades designability.

## G Broader Impact

NuCaliby’s broader impact lies in extending protein design from a residue-level problem to a joint optimization problem across the central dogma of molecular biology, enabling simultaneous control over protein structure, coding-sequence properties, and cellular expression context. This could improve the reliability and cost-efficiency of producing therapeutic proteins, vaccines, enzymes, and other biologics by directly incorporating objectives such as host-specific translation adaptation, mRNA stability, and regulatory RNA elements during sequence generation rather than through post-hoc optimization. The framework may also accelerate research in synthetic biology and biotechnology by enabling programmable integration of RNA-level functionality into designed proteins.

Like all protein-design technologies, NuCaliby carries a general dual-use risk because methods that improve biomolecular engineering could in principle be misapplied. However, the method does not provide a targeted capability for harmful applications beyond existing public protein-design tools. Its released models are comparable in capability to already-public residue-level inverse-folding approaches such as ProteinMPNN, CodonMPNN, and Caliby, and practical use still requires substantial domain expertise, experimental infrastructure, and wet-lab validation.

